# Molecular crowding facilitates ribozyme-catalyzed RNA assembly

**DOI:** 10.1101/2023.04.30.538884

**Authors:** Saurja DasGupta, Stephanie Zhang, Jack W. Szostak

## Abstract

Catalytic RNAs or ribozymes are considered to be central to primordial biology. Most ribozymes require moderate to high concentrations of divalent cations such as Mg^2+^ to fold into their catalytically competent structures and perform catalysis. However, undesirable effects of Mg^2+^ such as hydrolysis of reactive RNA building blocks and degradation of RNA structures are likely to undermine its beneficial roles in ribozyme catalysis. Further, prebiotic cell-like compartments bounded by fatty acid membranes are destabilized in the presence of Mg^2+^, making ribozyme function inside prebiotically relevant protocells a significant challenge. Therefore, we sought to identify conditions that would enable ribozymes to retain activity in low concentrations of Mg^2+^. Inspired by the ability of ribozymes to function inside crowded cellular environments with <1 mM free Mg^2+^, we tested molecular crowding as a potential mechanism to lower the Mg^2+^ concentration required for ribozyme-catalyzed RNA assembly. Here, we show that the ribozyme-catalyzed ligation of phosphorimidazolide RNA substrates is significantly enhanced in the presence of the artificial crowding agent polyethylene glycol. We also found that molecular crowding preserves ligase activity under denaturing conditions such as alkaline pH and the presence of urea. We also show that crowding-induced stimulation of RNA-catalyzed RNA assembly is not limited to phosphorimidazolide ligation but extends to the RNA-catalyzed polymerization of nucleoside triphosphates. RNA-catalyzed RNA ligation is also stimulated by the presence of prebiotically relevant small molecules such as ethylene glycol, ribose, and amino acids, consistent with a role for molecular crowding in primordial ribozyme function and more generally, in the emergence of RNA-based cellular life.

**Table of Contents Graphic:** 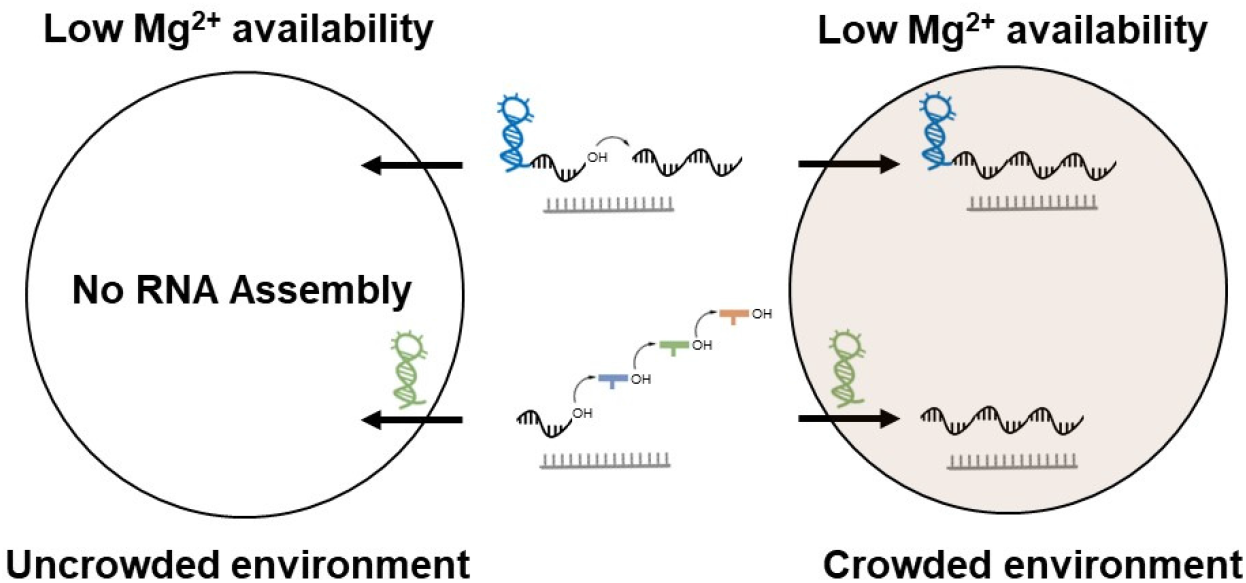

**SYNOPSIS:** Molecular crowding enables ribozyme-catalyzed RNA assembly under low Mg^2+^ conditions, and consequently may have played an essential role in the origin and evolution of RNA-based primordial life.

## INTRODUCTION

The catalytic repertoire of RNA lies at the foundation of the RNA world hypothesis, which posits that early life used RNA as both the genetic material and as enzymes (ribozymes).^1^ The ability of single-stranded RNA molecules to assume a wide range of folded structures endows them with functions such as molecular recognition and catalysis, suggesting that folded RNA structures would have been essential to early life. RNA assembly processes that could generate complex folded RNA structures were therefore likely to have played an important role in the propagation and evolution of the earliest living cells. Ribozymes usually require divalent cations like Mg^2+^ to access their functional folds and perform catalysis. Mg^2+^ facilitates RNA folding by partially neutralizing the negatively-charged RNA backbone, and often participates in catalytic interactions within the ribozyme active site.^2-4^ Although essential to RNA function, Mg^2+^ can also be detrimental. Mg^2+^ catalyzes RNA backbone hydrolysis, thereby disrupting functional structures. It also accelerates the hydrolysis of intrinsically reactive RNA building blocks such as phosphorimidazolides that would have been important for primordial RNA assembly.^5-7^ Additionally, Mg^2+^ is generally detrimental to the integrity of prebiotic cell-like compartments bounded by fatty acids, which are commonly used models of primordial cell membranes. This incompatibility between ribozyme function and the stability of protocell membranes poses a significant challenge toward efficient RNA catalysis within fatty acid protocells.^5^

RNA assembly would have driven primordial genetics and generated the catalytic diversity required to sustain RNA-based primordial life, therefore, ribozymes that catalyze RNA ligation or polymerization were crucial to primordial biology. Such ribozymes have been identified through *in vitro* evolution.^5^ Ligase and polymerase ribozymes that use 5′ triphosphorylated oligoribonucleotides and nucleoside triphosphates as substrates, respectively, exhibit high Mg^2+^ requirements. For example, the first-of-its-kind, class I ligase has a [Mg^2+^]_1/2_ of 70-100 mM^8^ and polymerase ribozymes derived from the class I ligase have an optimal [Mg^2+^] of ∼200 mM.^9, 10^ We previously reported ribozymes that catalyze ligation of RNA oligomers 5′ activated with a prebiotically plausible, 2-aminoimidazole (2AI) moiety. 2AI-activated RNA monomers/oligomers are useful substrates for nonenzymatic RNA assembly, therefore, these ‘2AI-ligase’ ribozymes provide continuity between chemical and enzymatic RNA ligation.^6^ Most of these 2AI-ligases were inefficient at low Mg^2+^ concentrations (<4 mM); however, we identified a single ligase sequence that had significantly lower Mg^2+^ requirement ([Mg^2+^]_1/2_ ∼0.9 mM).^11^ Although ribozymes with reduced Mg^2+^ requirements clearly exist, they are apparently relatively uncommon in RNA sequence space. We have therefore searched for a more general solution that would have enabled ribozymes to operate in low Mg^2+^ environments such as freshwater ponds or within protocells bounded by prebiotic fatty acids.^12^ Mechanisms that stimulate ribozyme activity at low [Mg^2+^] would lower the evolutionary threshold for the emergence of such molecules in the RNA world.

Although ribozymes usually require moderate to high Mg^2+^ concentrations to function *in vitro*, naturally-occurring ribozymes have evolved to function in the presence of 0.5-1 mM free Mg^2+^ within cellular environments.^13^ This lower Mg^2+^ requirement is thought to be a consequence of the crowded cellular environment. In addition to cellular structures like organelles, the intracellular milieu is crowded with molecules that range from biopolymers like nucleic acids and proteins to smaller molecules including amino acids, nucleotides, sugars, amines, and alcohols, which collectively occupy up to 30% of the cellular volume.^13, 14^ The presence of these molecules introduces a variety of physical and chemical forces that alter the properties of cellular RNAs.^15-17^ Volume excluded by macromolecules decreases the conformational entropy of unfolded RNA (an effect commonly referred to as ‘macromolecular crowding’) and consequently promotes RNA folding and RNA function. Unfavorable interactions between the solvent exposed RNA backbone and low MW species in the cellular milieu also induce folding to minimize these interactions. A decrease in dielectric constant may favor RNA-Mg^2+^ association due to the diminished solvation of free Mg^2+^, which can stimulate RNA folding and catalysis. A decrease in water activity caused by cosolutes may favor the formation of RNA folds with reduced solvent exposed surface area that is accompanied by water release.

Investigations into RNA structure and function in solutions artificially crowded with cosolutes like polyethylene glycol (PEG) have revealed favorable effects of crowding on RNA function.^17^ Biophysical studies using small angle x-ray scattering (SAXS) and single molecule Förster resonance energy transfer (smFRET) demonstrated that molecular crowding induces RNA folding. This effect is most pronounced in the low Mg^2+^ regime, where folded structures are not usually predominant.^18-20^ Enhanced folding in crowded solutions is often reflected in modest to significant increases in catalytic rates.^17^ Ribozymes in the RNA world may have evolved in similarly crowded environments either within primitive cellular compartments or confined microspaces on the Earth’s surface, which may have allowed them to function at low concentrations of Mg^2+^.^17, 21^

Here, we demonstrate the beneficial effects of molecular crowding on ribozyme-catalyzed RNA assembly, including the stimulation of ribozyme ligase activity at low millimolar concentrations of Mg^2+^ and the preservation of ribozyme activity under harsh reaction conditions such as alkaline pH or urea-induced denaturation. We propose that the stabilization of catalytic RNA folds in prebiotic crowded environments could provide a general means of enabling ribozyme-catalyzed RNA assembly in diverse environments including low [Mg^2+^] environments.

## RESULTS AND DISCUSSION

### Crowding rescues RNA-catalyzed RNA ligation at low Mg^2+^ concentrations

To test the effect of crowding on RNA-catalyzed RNA assembly, we chose a ligase ribozyme (henceforth, ligase 1) (Fig. 1A, Table S1), previously identified by *in vitro* selection, that catalyzes the template-directed ligation of a primer strand to a 2AI-activated oligonucleotide. This ribozyme exhibited significantly reduced product yield at Mg^2+^ concentrations below 4 mM.^6^ For example, ligation proceeded to ∼30% in 3 h at 4 mM Mg^2+^, but yields were reduced to 8%, 2%, and 1% at 3 mM, 2 mM, and 1 mM Mg^2+^, respectively. This ribozyme exhibits a corresponding reduction in activity in the low Mg^2+^ regime with only 5-15-fold rate enhancement over background at 2-3 mM Mg^2+^ compared to the ∼300-fold enhancement observed at 10 mM Mg^2+^.^6^ We used polyethylene glycol (PEG), to generate a crowded environment *in vitro*. PEG is chemically inert and available in a wide range of MWs, which allowed us to simulate the presence of a variety of small molecules and biopolymers that could have been present in prebiotic milieus. We also included ethylene glycol (EG) in our studies, in addition to PEGs of various MWs (PEG 200, PEG 400, PEG 1000, PEG 8000). EG can be synthesized abiotically^22, 23^ and is one the larger molecules detected in interstellar medium.^24, 25^ We first screened various concentrations of EG, PEG 200, PEG 400, PEG 1000, and PEG 8000 to identify optimal crowding conditions for ligase 1 activity in the presence of 1 mM Mg^2+^ and 100 mM Tris-HCl, pH 8 (Fig. S1). We observed remarkable ligation rescue in the presence of EG and both low and high MW PEGs. Ligation yield rose from barely detectable levels in the absence of crowding agents to about 20% and 50% after 3 h at 1 mM and 2 mM Mg^2+^, respectively, in the presence of 10% (*w/v*) EG. Similar stimulation in ligation was observed in 30% (*w/v*) PEG 200, PEG 400, 19% PEG 1000, 19% (*w/v*) PEG 8000 at 1 mM Mg^2+^ with ∼50% ligation after 3 h which is comparable to the 60% ligation observed in solution at 10 mM Mg^2+^ with no crowding agents (Fig. 1B, C). Ligation rates in the presence of crowders at low Mg^2+^ (0.7/h-1.3/h) were also comparable to the rate observed in the absence of crowders at 10 mM Mg^2+^ (∼1.5/h).

**Figure 1.**
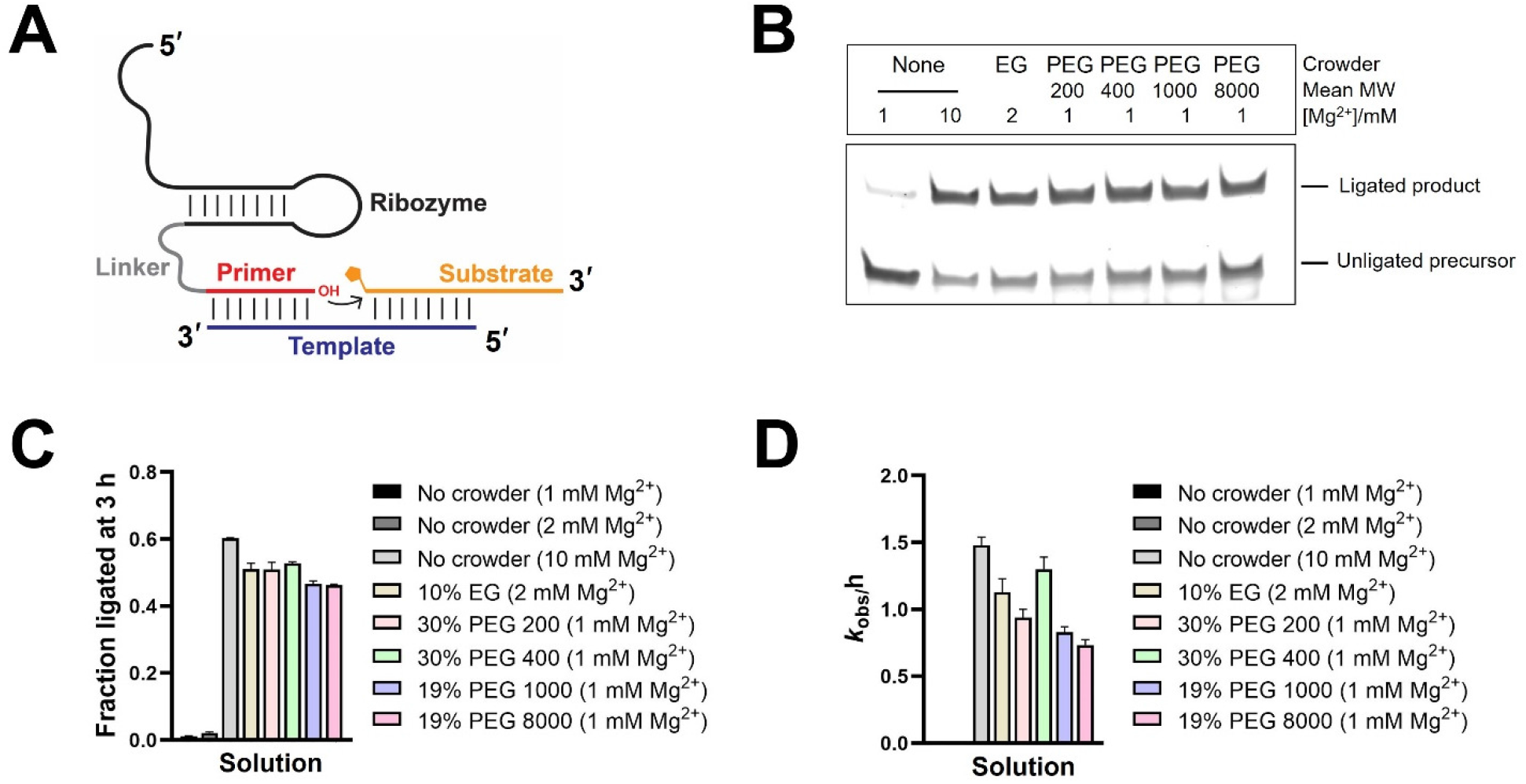
Stimulation of ribozyme activity of ligase 1 at 1-2 mM Mg^2+^ in the presence of ethylene glycol and PEGs. **A**. Schematic of ribozyme-catalyzed ligation of a 2-aminoimidazole-activated RNA substrate. **B**. Catalytic ligation is undetectable at 1-2 mM Mg^2+^ in solutions without crowders but is rescued in crowded solutions. **C**. Ligation yields after 3 h in the absence and presence of crowding agents at the indicated concentrations. **D**. Ligation rates in the absence and presence of crowding agents. Ligation reactions contained 1 µM ribozyme, 1.2 µM RNA template, and 2 µM 2-AI-activated RNA substrate in 100 mM Tris-HCl pH 8.0 and the indicated concentrations of MgCl_2_. Reactions contained additives (EG, PEG 200-8000) as indicated. None indicates the absence of crowders.

To understand the attenuated Mg^2+^ dependence of ribozyme-ligase activity, we measured ligation rates as a function of Mg^2+^ concentration in the presence of 10% EG (low MW additive), 30% PEG 200 (low MW additive), and 19% PEG 1000 (high MW additive). The concentration of Mg^2+^ at which half-maximum rate was achieved, [Mg^2+^]_1/2_, was significantly lowered in the presence of crowding agents (Fig. 2, Fig. S2), consistent with the enhanced ligation yield observed in a low Mg^2+^ concentration. A 3-fold reduction in [Mg^2+^]_1/2_ was observed in 10% EG while 30% PEG 200 and 19% PEG 1000 caused a ∼10-fold reduction (Fig. 2E). While all three crowders (EG, PEG 200, PEG 1000) supported ligase 1 activity at lower concentrations of Mg^2+^, maximal rates were achieved at sub millimolar Mg^2+^ with PEG 200 and PEG 1000, and at ∼2 mM Mg^2+^ with EG. Previous studies have found a decrease in Mg^2+^ requirement for ribozyme activity to accompany a decrease in Mg^2+^ requirement for folding, in both the group II intron^20^ and HDV^26^ ribozymes, supporting a role of crowding in facilitating the formation of catalytically relevant folds. An alternative explanation to the induction of RNA folding is that the addition of solutes like PEG may lower solution polarity and thereby result in greater association between RNA and Mg^2+^. An RNA-bound Mg^2+^ ion may activate the nucleophile at the site of ligation or stabilize the transition state.^27^ We tested nonenzymatic ligation in the presence of EG and various PEGs at 1 mM Mg^2+^ using a FAM-labeled primer (corresponding to the 3′ end sequence of the ligase downstream of the linker, Table S1), the 2AI-activated RNA substrate, and an appropriate RNA template. Ligation yields were unaffected in the presence of EG or PEGs (Fig. S3), supporting the importance of ribozyme structure in crowding-induced rate enhancement.

**Figure 2.**
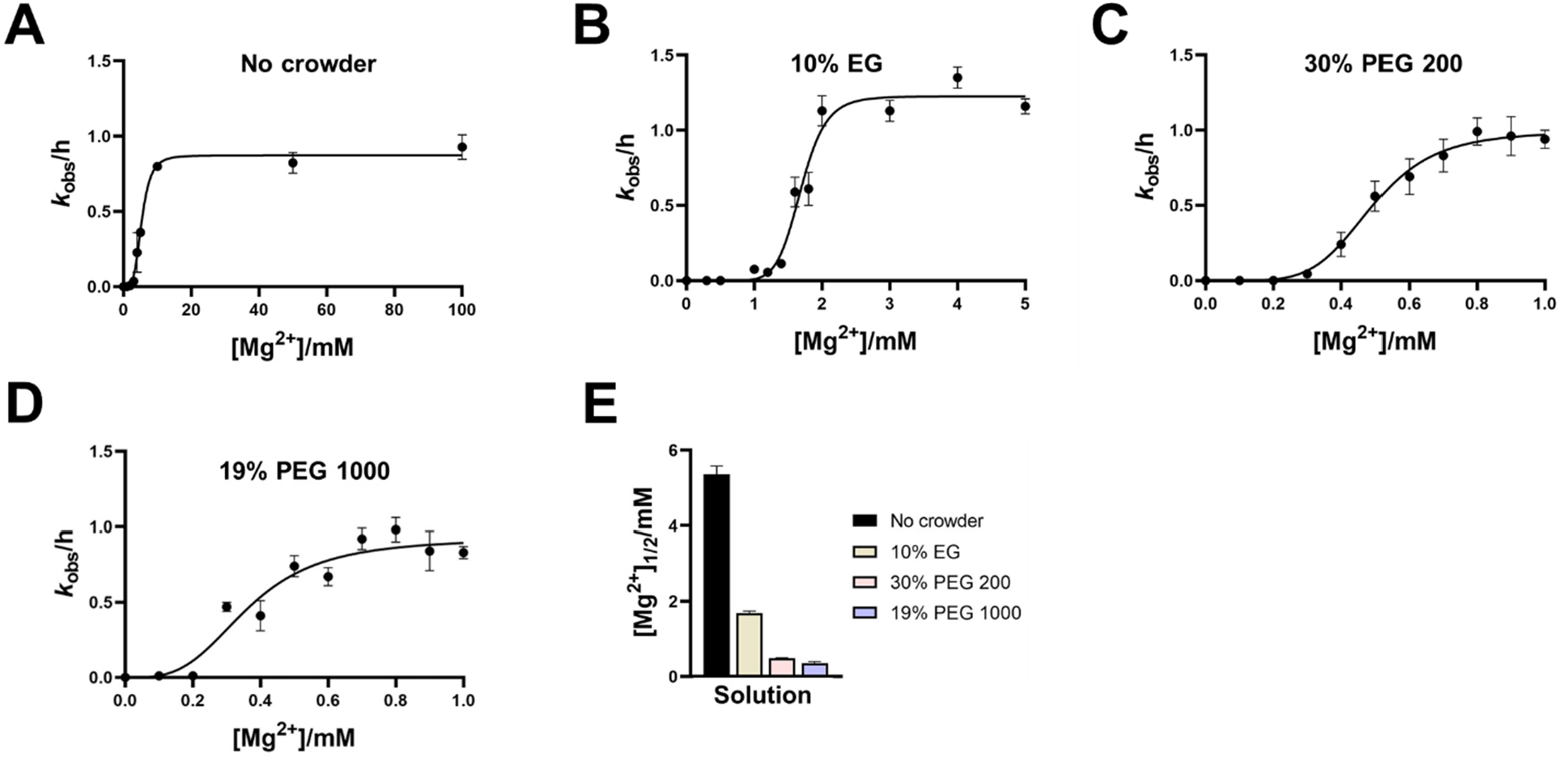
Crowding decreases the Mg^2+^ requirement for RNA-ligase activity. Mg^2+^ dependence on ligation rates of ligase 1 (A) in the absence of crowders, and in the presence of (B) 10% (*w/v*) EG, (C) 30% (*w/v*) PEG 200, and (D) 19% (*w/v*) PEG 1000. (E) Crowding agents reduce the [Mg^2+^]_1/2_ values for the rate of ribozyme ligation. Ligation reactions contained 1 µM ribozyme, 1.2 µM RNA template, and 2 µM 2-AI-activated RNA substrate in 100 mM Tris-HCl pH 8.0 and the indicated concentrations of MgCl_2_. Reactions contained additives (EG, PEG 200, or PEG 1000) as indicated.

To ask if the crowding stimulation of ribozyme-catalyzed phosphorimidazolide ligation was specific to ligase 1, or was more general, we tested the activity of another ligase ribozyme (henceforth, ligase 2) identified from our previous *in vitro* selection experiment (Table S1).^6^ Although distinct in sequence and structure, ligase 2 exhibited a similar response to crowding as ligase 1. While ∼6% ligation was observed in the absence of crowding agents after 3 h at 1 mM Mg^2+^, crowding increased ligation yields to ∼60%, which was comparable to the ligation yield at 10 mM Mg^2+^ in the absence of crowding agents. The rates of ligase 2-catalyzed ligation followed a similar trend (Fig. S4).

### Crowding protects ribozymes from denaturation

Since crowding promotes the formation of compact RNA folds, we wondered if molecular crowding could protect ribozymes from unfolding under denaturing conditions. Since ligase 1 and ligase 2 were inactive at low Mg^2+^ in the absence of crowding agents, we used a previously reported 2AI-ligase (henceforth, ligase 3) that is functional under these conditions for the following experiments.^11^ First, we tested RNA ligation by ligase 3 in the presence of urea, which is an effective denaturant of RNA and also an important precursor molecule in the prebiotic syntheses of RNA and amino acids.^28^ As expected, ligation rates in the background of 1 mM Mg^2+^ decreased by ∼6-fold in the presence of 1 M urea and ligation was further reduced in the presence of 2.5 M urea (Fig. 3A, Fig. S5A). Ligation in 1 M urea was restored upon addition of 30% PEG 200 and 19% PEG 1000 (Fig. 3A). Ligase 3 was even active in 2.5 M urea in the presence of PEG 200 and PEG 1000, with rate enhancements of 25-fold and 33-fold, respectively, relative to solutions without crowding agents. Interestingly EG did not show any rescue of ligation rate under these partially denaturing conditions (Fig. S5B). Ribozyme activity in the presence of molar concentrations of urea is consistent with the stabilization of compact, solvent-excluded RNA tertiary structures by crowding agents.^26^ We suggest that polymeric crowders such as polypeptides or polyesters or even ‘proto-peptides’ such as depsipeptides that contain a mixture of amide and ester linkages, if present in sufficient concentrations in prebiotic environments, could have shielded catalytic RNA structures from non-specific denaturation by molecules such as urea and formamide.^29^

**Figure 3.**
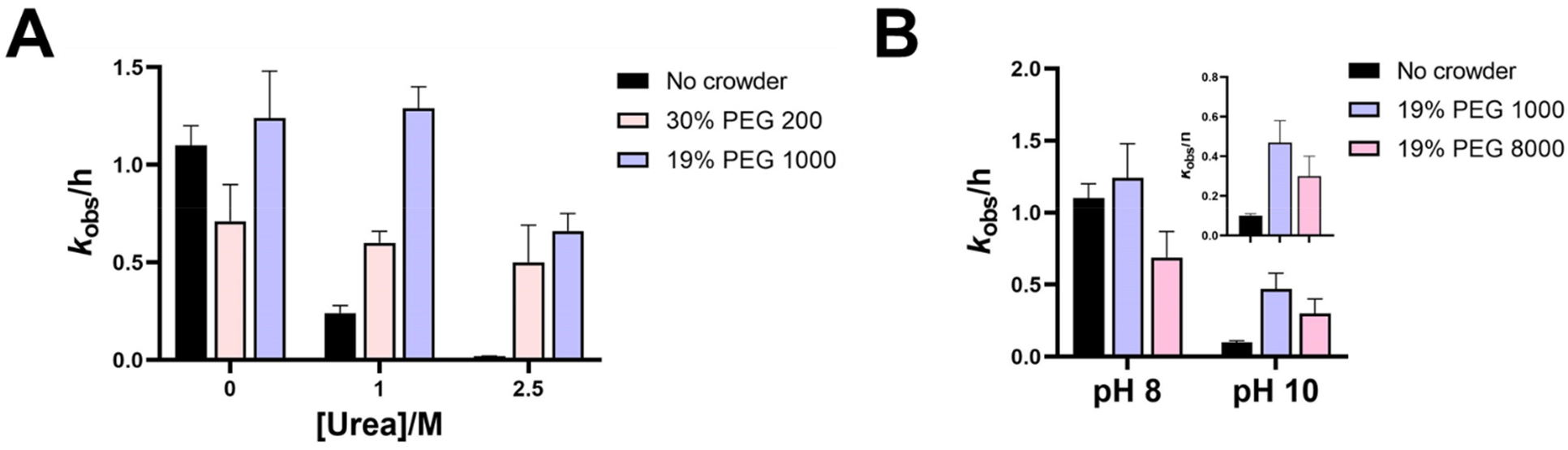
Molecular crowding counteracts loss of ribozyme ligase 3 activity under denaturing conditions. Crowding rescues the loss of ligation activity induced by (A) molar concentrations of urea and (B) alkaline pH. Ligation reactions contained 1 µM ribozyme, 1.2 µM RNA template, and 2 µM 2-AI-activated RNA substrate in 100 mM Tris-HCl pH 8.0, 1 mM MgCl_2_. Reactions contained additives (PEG 200, PEG 1000, or PEG 8000) and urea (1M or 2.5 M) as indicated.

Alkaline pH, which can be beneficial for certain prebiotic processes such as the synthesis of sugars^28^ and RNA strand separation,^30^ is detrimental to the chemical stability of RNA. However, compact folded RNAs are more resistant to alkaline degradation than their unfolded counterparts. Encouraged by the protective effect of crowding in the presence of urea, we measured the activity of ligase 3 at pH 10 and pH 11 in crowded solutions. No ligation was observed at pH 11 in the presence or absence of crowders. A small amount of ligated product was detected at pH 10 in the absence of crowding agents, with an 11-fold reduction in reaction rate relative to that in pH 8 (*k*_obs_ values of 1.1/h at pH 8 vs. 0.1/h at pH 10). The loss of ligase activity at pH 10 was less pronounced in the presence of PEG 1000 and PEG 8000 with only a 2.6-fold and 2.3-fold reduction in *k*_obs_, respectively relative to their values at pH 8 (Fig. 3B, Fig. S6A). This represents a 4-5-fold rate enhancement for ribozyme-catalyzed ligation at pH 10 in crowded solutions (Fig. 3B). Although ligase activity was rescued in the presence of crowders, crowding had minimal effect on the extent of RNA degradation at pH 10 or pH 11. Therefore, the beneficial effect of crowders may result from the protection of the catalytic fold from disruption at alkaline pH, or by preserving base-pairing interactions between the substrate, template, and ribozyme.

We asked whether crowding could have had a similar protective function during fluctuating temperature cycles on the early Earth. However, only a modest enhancement in ligation rates by ligase 3 was observed at 55 ºC in the presence of EG, PEG 400, PEG 1000, and PEG 8000 (Fig. S6B, C). The lack of substantial benefit from crowding at high temperatures is consistent with UV melting experiments with ligase 1 ribozyme, which revealed a negligible increase in its thermal stability in the presence of both low (EG) or high (PEG 1000) MW crowders (Fig. S7A, B).

### Crowding stimulates RNA-catalyzed RNA polymerization at low Mg^2+^ concentrations

Although ribozymes that catalyze the template-directed polymerization of nucleoside phosphorimidazolides have not yet been reported, polymerase ribozymes that use NTPs as substrates have been evolved from the class I ligase ribozyme.^9, 10, 31^ These ribozymes generally require 50-200 mM Mg^2+^ which makes them incompatible with fatty acid vesicle-based models for primitive cells. Tagami *et al*. demonstrated modest polymerase function at 10 mM free Mg^2+^ in the presence of lysine decapeptide (K_10_), which enabled RNA-catalyzed RNA polymerization within Mg^2+^ resistant 1-palmitoyl-2-oleoylphosphatidylcholine (POPC) vesicles.^32^ We wondered if PEG could substantially lower the Mg^2+^ requirement of polymerase ribozymes in a manner similar to the 2AI-ligase ribozymes. We tested the ability of the 38-6 polymerase ribozyme^10^ to extend a 10 nt RNA primer on a 21 nt RNA template in the presence of 5 mM Mg^2+^ in solutions containing low or high MW PEGs. Negligible extension beyond +4 was observed in the absence of crowding agents; however, small amounts of full-length products (+11) were detected in the presence of PEG 200 or PEG 1000 after 1 day. The prominent +1 extension product increased from 24% without crowding agents to 33% and 40% in the presence of PEG 200 and PEG 1000, respectively. While only 26% of the primer was extended in the absence of crowding agents, 37% and 43% of the primer was extended in the presence of PEG 200 or PEG 1000, respectively (Fig. S7). This enhancement of ribozyme polymerase activity at low millimolar Mg^2+^ underscores the generality of the beneficial effects of crowded environments on ribozyme-catalyzed RNA assembly. Interestingly, molecular crowding has also been found to enhance the polymerization of NTPs^33^ and dNTPs^34^ by biologically-derived protein polymerases, which further supports the role of crowding in facilitating nucleic acid assembly.

### Prebiotically relevant small molecules enable ribozyme-catalyzed RNA ligation at low Mg^2+^

While our observations on the effects of molecular crowding agents on ribozyme activity are promising, the above results were obtained with prebiotically-irrelevant synthetic PEG molecules, except for EG. Therefore, we explored the potential of prebiotically-relevant small molecules for stimulating ribozyme-ligase activity. Considering the importance of simple sugars in a pre-RNA/RNA world, and the stabilizing effect of ribose on fatty acid membranes, we decided to explore the effect of ribose on ligase 1 ribozyme activity. ^28 35, 36^ We also tested a subset of amino acids thought to be available on early earth as products of prebiotic synthetic pathways such as the cyanosulfidic protometabolic reaction network.^37^ Ribose at 2% (*w/v*) and 3.8% (*w/v*) significantly increased ribozyme ligation yield from ∼1% ligation to ∼11% and ∼26% ligation after 3 h in the presence of 1 mM Mg^2+^ with *k*_obs_ values of ∼0.4/h and ∼0.5/h, respectively (Fig. S9). We also screened the amino acids glycine, alanine, proline, leucine, serine, and aspartic acid at 2.5 mM, 5 mM, 10 mM, and 20 mM concentrations for their ability to stimulate ligation at 1 mM Mg^2+^. All the above amino acids were found to stimulate ligation regardless of their concentrations with yields of 25%-35% after 3 h (Fig. S10). As lower concentrations are prebiotically more likely in most microenvironments, we measured the yield and rate of ligase 1-catalyzed RNA ligation in presence of 2.5 mM of each amino acid and a mixture of all six amino acids at a total concentration of 2.5 mM. The presence of amino acids both individually and as a mixture rescued ligation rates to within a factor of 2-3 of that observed at 10 mM Mg^2+^ without any additive (Fig. 4). This ability of prebiotic small molecules to facilitate ribozyme-catalyzed RNA assembly presents a ‘systems’ level solution for lowering the Mg^2+^ requirement for this central process in primordial biochemistry.

**Figure 4.**
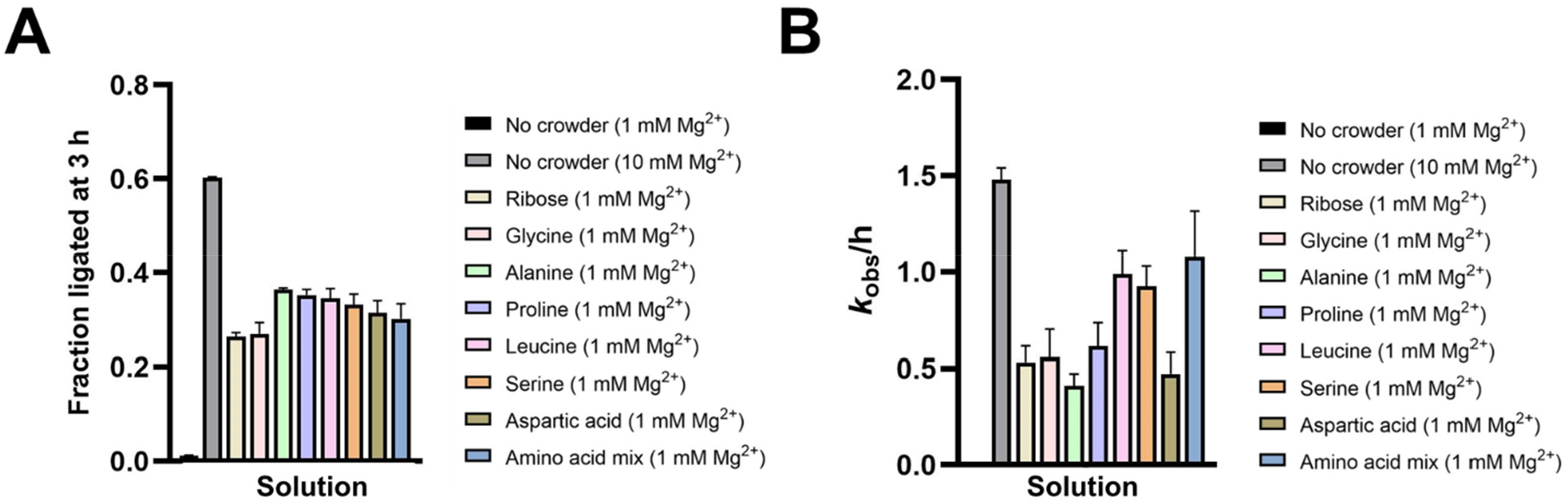
RNA ligation catalyzed by ligase 1 ribozyme is stimulated in the presence of prebiotic molecules. **A**. Ligation yields after 3 h at 1 mM Mg^2+^ in the presence of ribose and prebiotic amino acids. **B**. Ligation rates at 1 mM Mg^2+^ in the presence of ribose and prebiotic amino acids. Ligation reactions contained 1 µM ribozyme, 1.2 µM RNA template, and 2 µM 2-AI-activated RNA substrate in 100 mM Tris-HCl pH 8.0 and 1 mM or 10 mM MgCl_2_. Reactions contained additives (ribose, individual amino acid, or amino acid mixture) as indicated.

## CONCLUSION

The crucial role of Mg^2+^ in both nonenzymatic and ribozyme-catalyzed RNA replication, coupled with its ability to accelerate RNA degradation and destabilize fatty acid protocells, presents a puzzle for the emergence of RNA-based cellular life. Therefore, exploring scenarios that mitigate this ‘Mg^2+^ problem’ is of critical importance. The low Mg^2+^ requirement for natural ribozymes that function within crowded cellular environments inspired us to study molecular crowding as a general solution to the Mg^2+^ problem in the context of ribozyme-catalyzed RNA assembly. Our results show a dramatic stimulation of ribozyme-catalyzed ligation of 2AI-activated RNA oligomers at low millimolar Mg^2+^ by prebiotically-relevant amino acids, ribose, ethylene glycol, and polyethylene glycols of various MWs (200-8000). The beneficial effects of amino acids, ribose and ethylene glycol are especially notable since these molecules can be synthesized abiotically and therefore were likely to have been present in early Earth environments. The 3-10-fold lower Mg^2+^ requirement for ligase ribozymes in the presence of such solutes likely stems from enhanced RNA folding in ‘crowded’ solutions as the corresponding nonenzymatic ligation reaction was not affected by crowding. Stimulation of catalytic activity in the presence of molecular crowding has been reported for other ribozymes.^17^ Since the crowding-induced enhancement of ligation was largely independent of crowder size, ribozyme folding could be favored by an interplay of both enthalpic forces arising from interactions between the RNA surface and the crowder and entropic forces arising from volume exclusion.^17^ Further studies may help isolate the effects of these distinct thermodynamic forces.

We demonstrated that in addition to enabling RNA ligation in the low Mg^2+^ regime, crowding offers modest to significant protection to ligase ribozymes under various denaturing conditions relevant to early earth environments. The ability to function under conditions that favor the disruption of RNA secondary structure could have been important for rapid RNA-catalyzed RNA replication which requires the separation of newly synthesized RNA strands from their RNA templates while preserving catalytic RNA structures. The beneficial effects of crowding are not limited to 2AI-ligase ribozymes but extend to RNA-catalyzed polymerization of NTPs, a reaction that has a significantly higher Mg^2+^ requirement than the ligation of phosphorimidazolide RNA substrates.

Efficient RNA assembly, at low Mg^2+^ concentrations, presents a path to reconcile ribozyme function with the stability of protocell membranes made of fatty acids. Protocells crowded with prebiotic small molecules like sugars, alcohols, and amines, and polymeric species such as short oligonucleotides or polypeptides could potentially support a wide range of ribozyme activities under the low Mg^2+^ concentrations required for maintaining membrane integrity. This is particularly interesting in the context of our earlier observation that prebiotically relevant small molecules including ribose also reduce RNA leakage from fatty acid vesicles.^11^ The combined effect of enhancing ribozyme function under low Mg^2+^ conditions and stabilizing protocell membranes against Mg^2+^ suggests a potential role for these prebiotic molecules that is separate from their roles as components of the building blocks of life. By providing a general mechanism to activate RNA catalysis at low Mg^2+^ concentration, molecular crowding expands the range of environments in which ribozymes can function to less salty environments such as freshwater bodies^12^ and increases the likelihood of the emergence of active ribozymes from RNA sequence space.^21^ Suboptimal sequences that would otherwise not be selected in low Mg^2+^ environments could emerge in crowded milieus, potentially creating neutral mutational pathways that would facilitate ribozyme evolution and therefore increase the catalytic diversity of the RNA world.

## EXPERIMENTAL PROCEDURES

### RNA preparation and substrate activation

Ribozymes were prepared by *in vitro* transcription of PCR-generated dsDNA templates containing 2′-*O*-methyl modifications to reduce transcriptional heterogeneity at the 3′ end of the RNA^38^ (Table S1). Transcription reactions contained 40 mM Tris-HCl (pH 8), 2 mM spermidine, 10 mM NaCl, 25 mM MgCl_2_, 10 mM dithiothreitol (DTT), 30 U/mL RNase inhibitor murine (NEB), 2.5 U/mL thermostable inorganic pyrophosphatase (TIPPase) (NEB), 4 mM of each NTP, 30 pmol/mL DNA template, and 1U/µL T7 RNA Polymerase (NEB) and were incubated for 3 h at 37°C. DNA template was digested by DNase I (NEB) treatment and RNA was extracted with phenol-chloroform-isoamyl alcohol (PCI), ethanol precipitated, and purified by denaturing PAGE. Ligation templates, FAM-labeled primers, and ssDNA were purchased from Integrated DNA Technologies.

The 5′ monophosphorylated oligonucleotide corresponding to the substrate sequence was activated by incubating it with 0.2 M 1-ethyl-3-(3 dimethylaminopropyl) carbodiimide (HCl salt) and 0.6 M 2-aminoimidazole (HCl salt, pH adjusted to 6) for 2 h at room temperature. The reaction was washed with water in Amicon Ultra spin columns (3kDa cutoff) four-five times (200 µL water per wash) and purified by reverse phase analytical HPLC using a gradient of 98% to 75% 20 mM TEAB (triethylamine bicarbonate, pH 8) versus acetonitrile over 40 min.^6^

### Ligation assays

Ligation reactions contained 1 µM ribozyme, 1.2 µM RNA template, and 2 µM 2-AI-activated RNA substrate in 100 mM Tris-HCl pH 8.0, the indicated concentrations of MgCl_2,_ and crowding agents. Aliquots were quenched with 5 volumes of quench buffer (8M urea, 100 mM Tris-Cl, 100 mM boric acid, 100 mM EDTA) and analyzed by denaturing PAGE. Gels were stained using SYBR™ Gold, imaged on an Amersham Typhoon RGB instrument (GE Healthcare), and analyzed in ImageQuant IQTL 8.1. Kinetic data were nonlinearly fitted to the modified first order rate equation, y = A (1 – e^-*k*x^), where A represents the fraction of active complex, *k* is the first order rate constant, x is time, and y is the fraction of ligated product in GraphPad Prism 9. For nonenzymatic ligation, a 5′ FAM-labeled RNA primer corresponding to the last 8 nt of the ribozyme sequence was used instead of the ribozyme, and the gel was directly imaged.

#### Ligation assays under denaturing conditions

All ligation assays under denaturing conditions were performed with ligase 3 ribozyme, which retains activity under low [Mg^2+^].

##### Ligation at high pH

ribozyme and template were heated at 95 °C for 2 min in the absence of any buffer and cooled to room temperature. CAPS buffer (pH 10 or pH 11) was added to a final concentration of 100 mM in the absence or presence of crowding agents (19% PEG 1000 or 19% PEG 8000) and 1 mM MgCl_2_. The substrate was added immediately after the addition of MgCl_2_ to initiate ligation.

##### Ligation at high temperatures

reactions with or without crowding agents (10% ethylene glycol or 30% PEG 400 or 19% PEG 1000 or 19% PEG 8000) were incubated at 55 °C after initiating ligation by adding the substrate.

##### Ligation in presence of urea

10 M urea was added to final concentrations of 1 M or 2.5 M after refolding in the presence of crowding agents (30% PEG 200 or 19% PEG 1000) and 1 mM MgCl_2_ to minimize degradation at high temperatures required for refolding. The substrate was added immediately after the addition of MgCl_2_ to initiate ligation.

### Ribozyme-catalyzed NTP polymerization assays

A FAM-labeled RNA primer (80 nM), RNA template (100 nM), and polymerase ribozyme (100 nM) were heated in the absence and presence of crowding agents and 25 mM Tris·HCl pH 8 at 80 °C for 30 s and cooled to 17 °C at a gradient of 0.1 °C/s. MgCl_2_ was added to final concentrations of 5 mM or 200 mM, followed by 0.5 mM of each NTP. Reactions were incubated at 17 °C for 1 day and 1 µL aliquots were quenched with 7 µL quench buffer (8M urea, 100 mM Tris-Cl, 100 mM boric acid, 100 mM EDTA containing 5 µM DNA oligo complementary to template). Reactions were analyzed by denaturing PAGE. Gels were imaged on an Amersham Typhoon RGB instrument (GE Healthcare) and analyzed in ImageQuant IQTL 8.1.

### UV melting analysis of ligase ribozyme

UV melting experiments were performed to determine the thermal stability of ligase 1 ribozyme in the absence of presence of low and high MW crowding agents according to the protocol used by Struslon *et al*.^39^. Briefly, 0.5 µM ribozyme was incubated at 95 °C for 2 min in 10 mM sodium cacodylate buffer (pH 7), and refolded in the absence or presence of crowding agents (10% ethylene glycol or 19% PEG 1000) in presence of 1 mM MgCl_2_ by heating the solution to 55 °C for 10 min followed by cooling to room temperature for 10 min. A Cary UV-Vis multicell peltier spectrophotometer was used for melting experiments. Absorbance was recorded at 260 nm every minute between 20 °C and 90 °C. Data was normalized with respect to ‘buffer only’ sample in each case, which contained all components in the experimental sample except RNA. Derivative plots of normalized data (dA/dT) vs T) and melting temperatures (T_*m*_) were obtained by the instrument’s default software.

## Acknowledgment

J.W.S. is an Investigator of the Howard Hughes Medical Institute. This work was supported in part by a grant from the Simons Foundation (290363) to J.W.S.

