## Supplemental Information for "Molecular crowding facilitates ribozyme-catalyzed RNA assembly"

### This PDF file includes:

Table S1
Figures S1 to S10

| Name | Sequence (5′→3′) |
|---|---|
| Ligase 1 | GACUCACUGACACAGAUCCACUCACGGACAGCGAAGCUCUCGCCAGCAAAAGAACAGACCGUCGAGGAAACGGCGCUGUCCUUUUUUGGCUAAG**G** |
| Ligase 2 | GACUCACUGACACAGAUCCACUCACGGACAGCGGAAUGCUGCCAACCGUGCGGGCUAAUUGGCAGACUGAGCUCGCUGUCCUUUUUUGGCUAAG**G** |
| Ligase 3 | GACUCACUGACACAGAUCCACUCACGGACAGCGAGCCACUGCGGAAGACCUUAAGAGGUGUAAUUGCUCACCCCGCUGUCCUUUUUUGGCUAAG**G** |
| Ligation primer | GGCUAAG**G** |
| Ligation template | GCGGUGGUCCUUAGCC |
| 2AI-activated substrate | (5′-phosphoro-2AI)-ACCACCGCAUUCCGCA |
| 38-6 polymerase | G*GUCAUUG*CCGCACAAAGACAAAUCUCCCCUCAGAGCUUGAGAACAUCUACGGAUGCAGAGGAGGGGCCUUCGGUGGAUCAAUUGUGCACCACCGUUCUCAACACGUACCCGAACAUAAAAAGACCUGACAAAAAGGCGAUGUUAGACACGCACAGGUGCCAUACCCAACACAUGGCUGAC |
| Polymerization primer | CACUCCACA**C** |
| Polymerization template | GAC*AAUGAC*AAAAAAAUCAGUACGUCGUGUGGAGUG |

**Table S1. RNA oligonucleotides used in this study.** Fixed base-paired stem nucleotides are underlined. Primer sequences are shown in red and the nucleotide containing the 3′ OH nucleophile is boldfaced. Template sequences complementary to primers are shown in orange. Sequences in 38-6 polymerase and the polymerization template that are complementary and contribute to ribozyme-template binding are *italicized.*

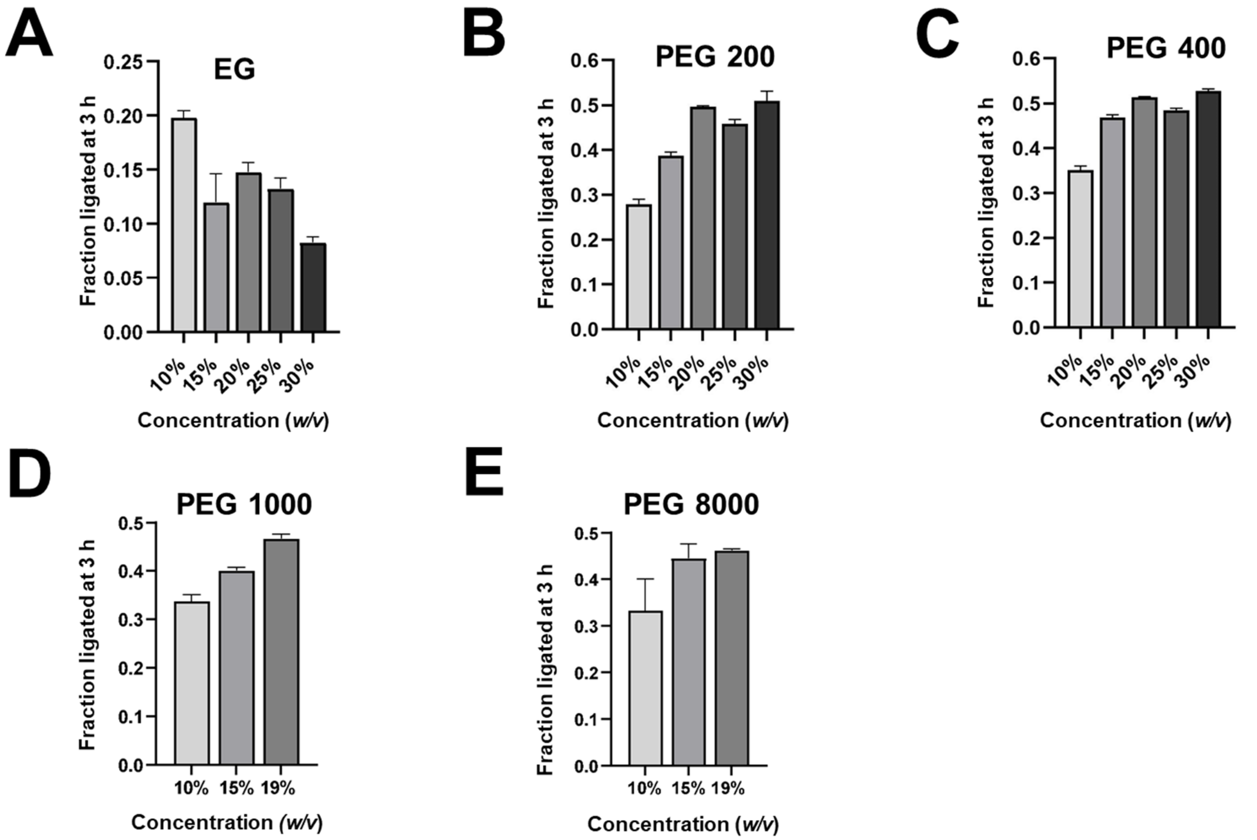


**Figure S1. Identification of optimal crowding conditions for RNA-catalyzed RNA ligation by ligase 1.** Fraction ligated after 3 h in the presence of (A) 10-30% ethylene glycol (EG), (B) 10-30% polyethylene glycol (PEG) 200, (C) 10-30% PEG 400, (D) 10-19% PEG 1000, (E) 10-19% PEG 8000. Ligation reactions contained 1 µM ribozyme, 1.2 µM RNA template, and 2 µM 2-AI-activated RNA substrate in 100 mM Tris-HCl pH 8.0 and 1 mM $MgCl_2$. Reactions contained additives (EG, PEG 200-8000) as indicated.

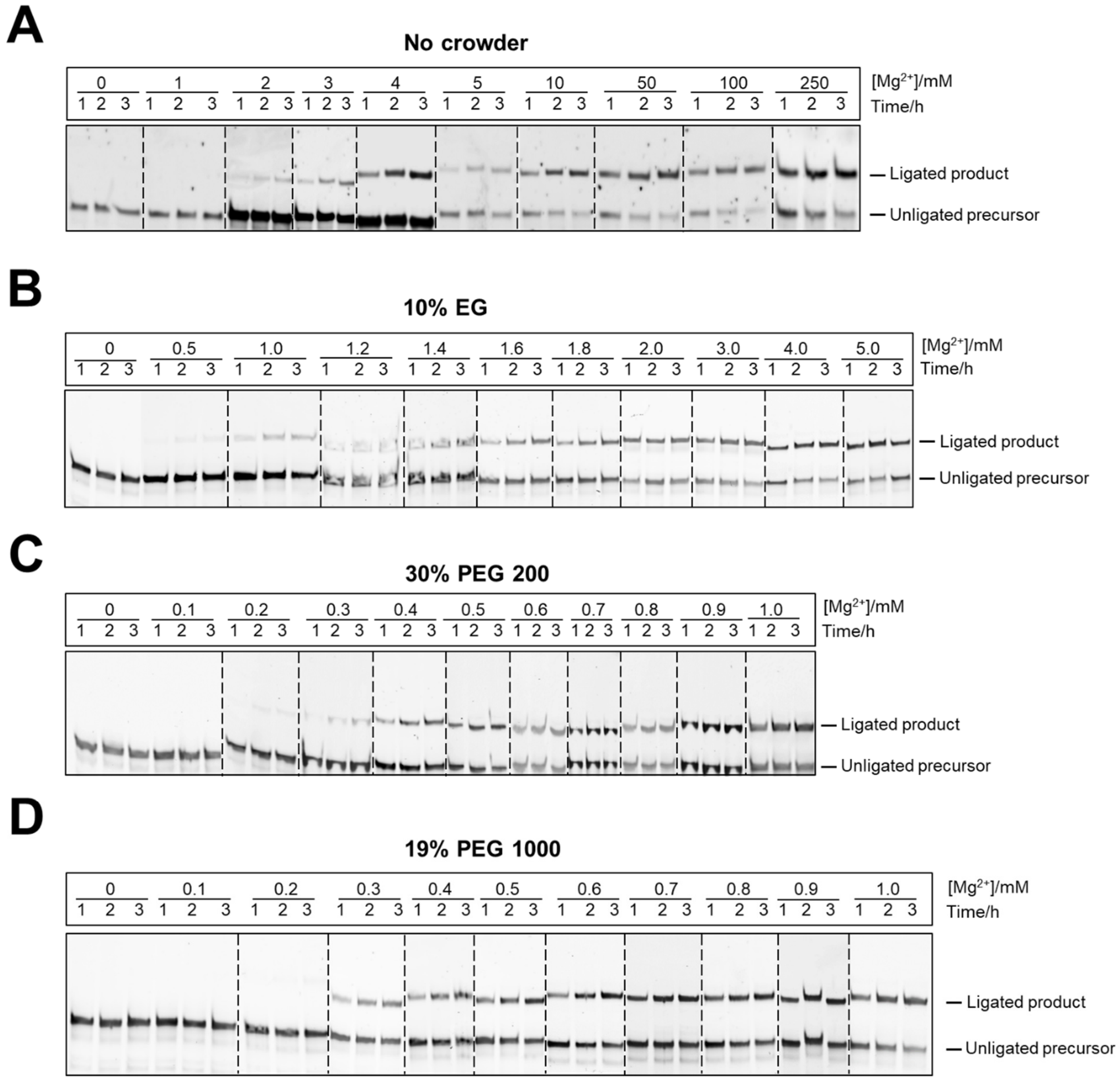


**Figure S2. Representative gels illustrating $Mg^{2+}$ dependence of ligase 1 ribozyme-catalyzed ligation in the absence and presence of crowding agents. A.** No crowding agents. **B.** 10% EG. **C.** 30% PEG 200 **D.** 19% PEG 1000. Ligation reactions contained 1 µM ribozyme, 1.2 µM RNA template, and 2 µM 2-AI-activated RNA substrate in 100 mM Tris-HCl pH 8.0 and the indicated concentrations of $MgCl_2$. Reactions contained additives (10% EG, 30% PEG 200, or 19% PEG 1000) as indicated.

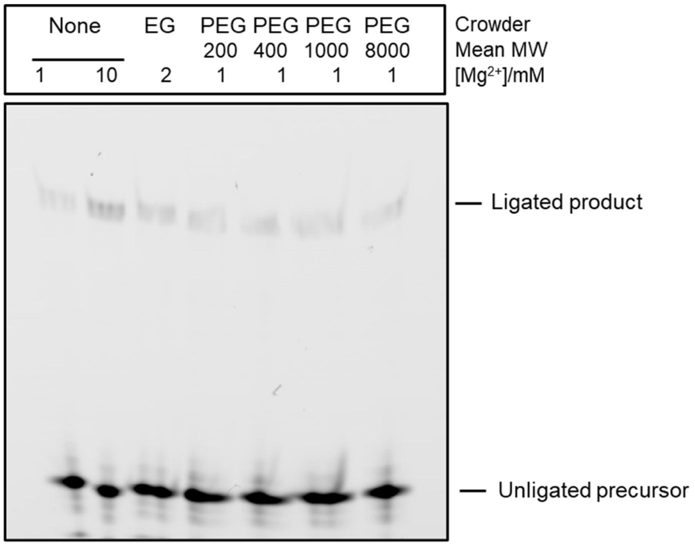


**Figure S3. Nonenzymatic ligation is not influenced by molecular crowding.** EG or PEGs do not rescue template-directed nonenzymatic RNA ligation at low $Mg^{2+}$ concentrations. Ligation reactions contained 1 µM FAM-labeled RNA primer, 1.2 µM RNA template, and 2 µM 2-AI-activated RNA substrate in 100 mM Tris-HCl pH 8.0 and 1 mM, 2 mM, or 10 mM $MgCl_2$. Reactions contained additives (EG, PEG 200-8000) as indicated. None indicates the absence of crowders.

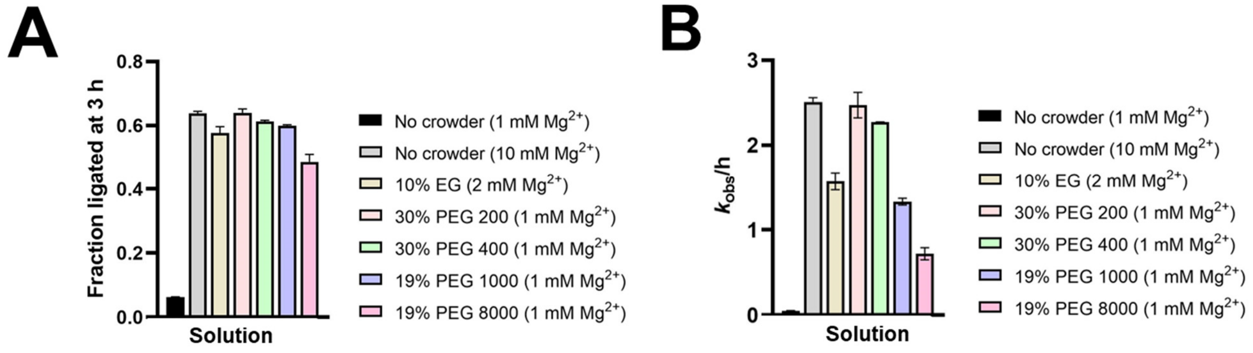


**Figure S4. Ligase 2 activity at low $Mg^{2+}$ concentrations is rescued by crowding agents.** In the presence of 1-2 mM $Mg^{2+}$, crowding agents stimulate (A) ligation yields and (B) ligation rates. Ligation reactions contained 1 µM ribozyme, 1.2 µM RNA template, and 2 µM 2-AI-activated RNA substrate in 100 mM Tris-HCl pH 8.0 and the indicated concentrations of $MgCl_2$ (1 mM, 2 mM, or 10 mM). Reactions contained additives (EG, PEG 200-8000) as indicated.

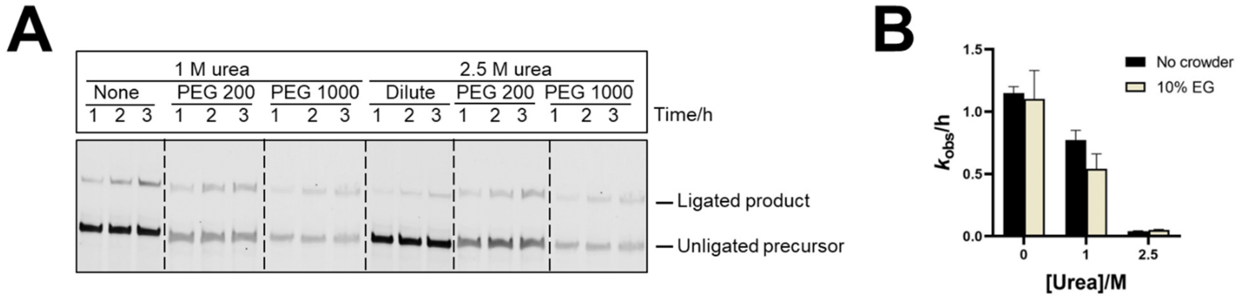


**Figure S5. Effect of molecular crowding on ligase 3-catalyzed RNA ligation in the presence of urea. A.** Ligase 3 activity in the presence of urea in the absence or presence of crowding agents. **B.** EG does not impart any beneficial effect to ligation in the presence of urea. Ligation reactions contained 1 µM ribozyme, 1.2 µM RNA template, and 2 µM 2-AI-activated RNA substrate in 100 mM Tris-HCl, pH 8.0, 1mM (A) or 2 mM (B) $MgCl_2$ in the presence or absence of urea. Reactions contained additives as indicated. None indicates the absence of crowding agents.

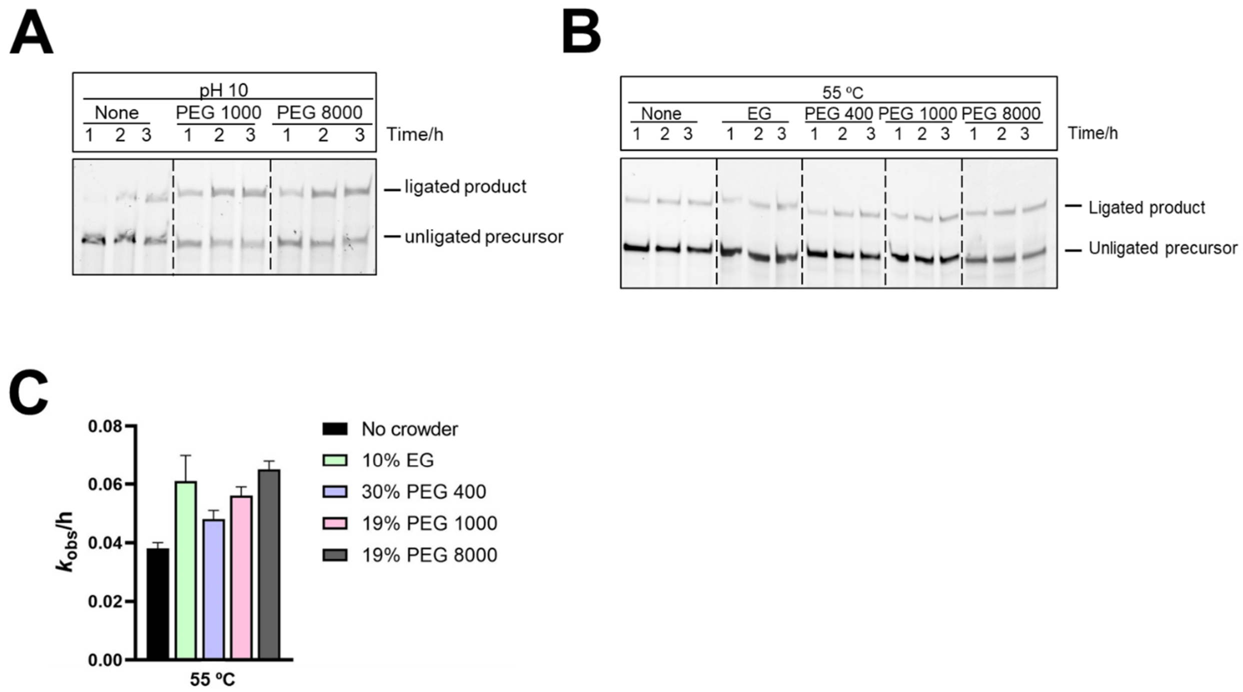


**Figure S6. Effect of molecular crowding on ligation by ligase 3 under alkaline pH and high temperature. A.** Ligase 3 activity at pH 10 in the absence or presence of crowding agents. **B.** Ligase 3 activity at 55 ºC in the absence or presence of crowding agents. **C.** Ligase 3 exhibits a modest increase in ligation rates upon addition of EG and PEG at 55 °C in 1 mM $Mg^{2+}$. Ligation reactions contained 1 µM ribozyme, 1.2 µM RNA template, and 2 µM 2-AI-activated RNA substrate in 100 Mm buffer: CAPS pH 10 (A) or Tris-HCl pH 8.0 (B). Reactions contained additives (EG, PEG 200-8000) as indicated. None indicates the absence of crowding agents.

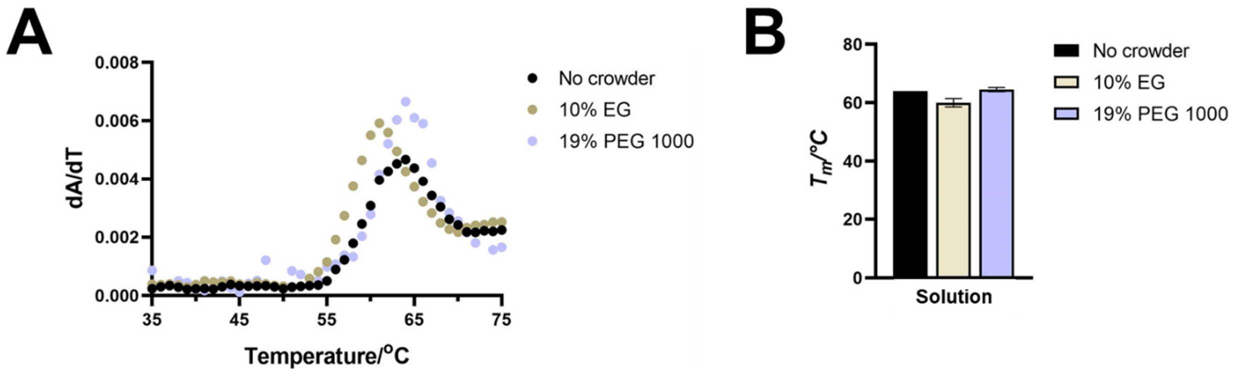


**Figure S7. Effect of molecular crowding on the thermal stability of ligase 1 ribozyme.** Thermal denaturation of ligase 1 measured by UV spectroscopy in 1 mM $Mg^{2+}$. $(dA/dT)_{max}$ occurs around the same temperature in both uncrowded and crowded solutions. **B.** Melting temperature (*T*m) of ligase 1 in uncrowded solution was comparable to the *T*m values observed in the presence of EG or PEG 1000. Thermal denaturation was performed with 0.5 µM ribozyme in 10 mM sodium cacodylate buffer, pH 7.

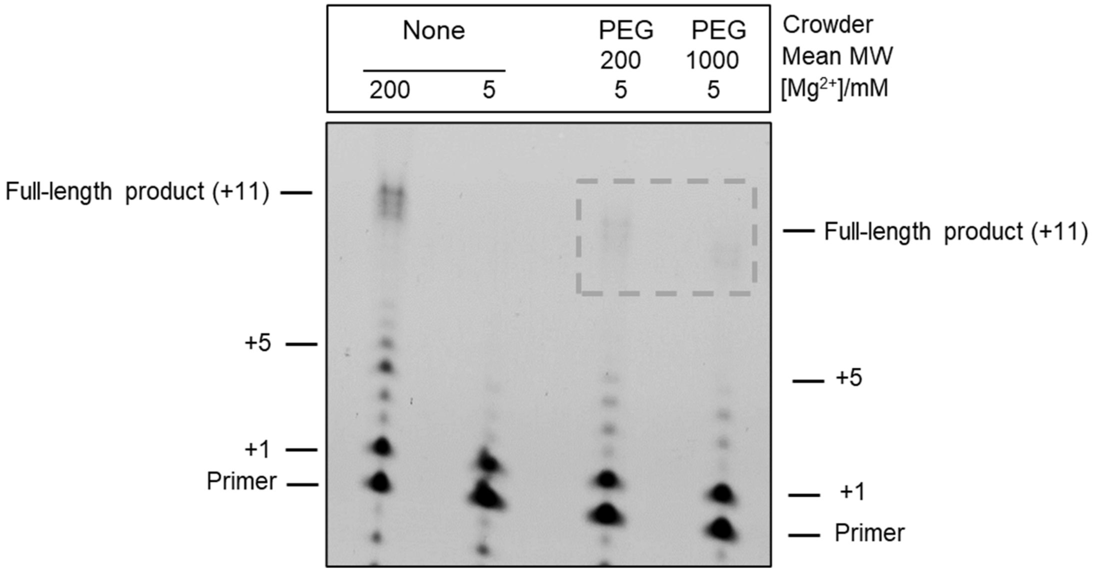


**Figure S8. RNA-catalyzed polymerization of NTPs at low $Mg^{2+}$ concentration.** Full length ribozyme (38-6 polymerase)-catalyzed polymerization product was detected at 5 mM $Mg^{2+}$ in reactions containing PEG 200 and PEG 1000; however, no such product was detected after 1 day in an uncrowded solution containing 5 mM $Mg^{2+}$. At 5 mM $Mg^{2+}$, crowders increased the fraction of extended primer from 26% in uncrowded solution to ~40%. Polymerization reactions containing a FAM-labeled RNA primer (80 nM), RNA template (100 nM), 2 mM total NTPs, and polymerase ribozyme (100 nM) in 25 mM Tris-HCl, pH 8 and 5 mM or 200 mM $Mg^{2+}$ were incubated at 17 °C. None indicates the absence of crowders.

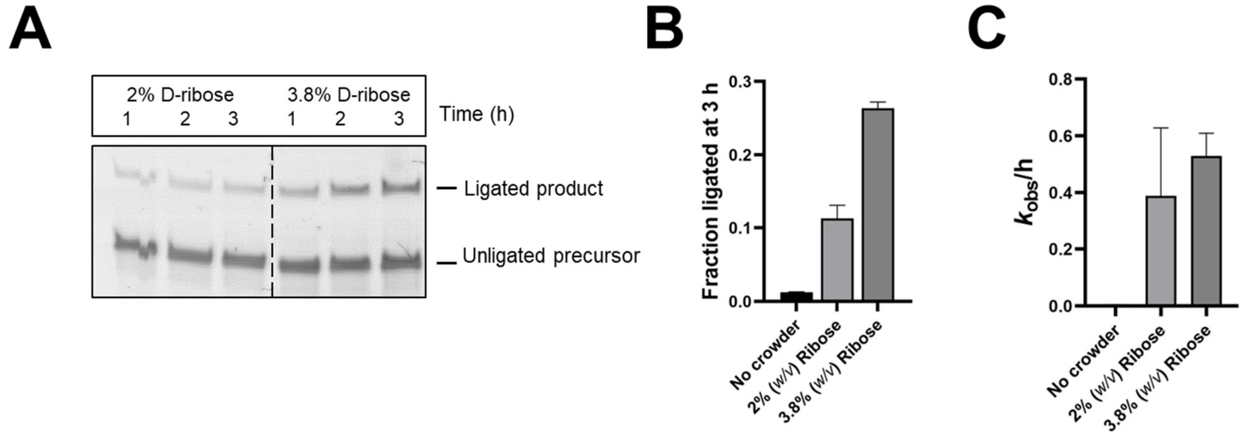


**Figure S9. Ribozyme-catalyzed RNA ligation is stimulated in the presence of ribose. A.** Ligase 1 ribozyme is active at 1 mM $Mg^{2+}$ in the presence of 2% and 3.8% D-ribose. **B.** Ligation yields after 3 h at 1 mM $Mg^{2+}$ increased to 11% and 26% from ~1% in the presence of 2% and 3.8% ribose, respectively. **C.** Ligation rates were enhanced significantly in the presence of 2% and 3.8% ribose. Ligation reactions contained 1 µM ribozyme, 1.2 µM RNA template, and 2 µM 2-AI-activated RNA substrate in 100 mM Tris-HCl pH 8.0 and 1 mM $MgCl_2$. Reactions were performed in the absence or presence of D-ribose.

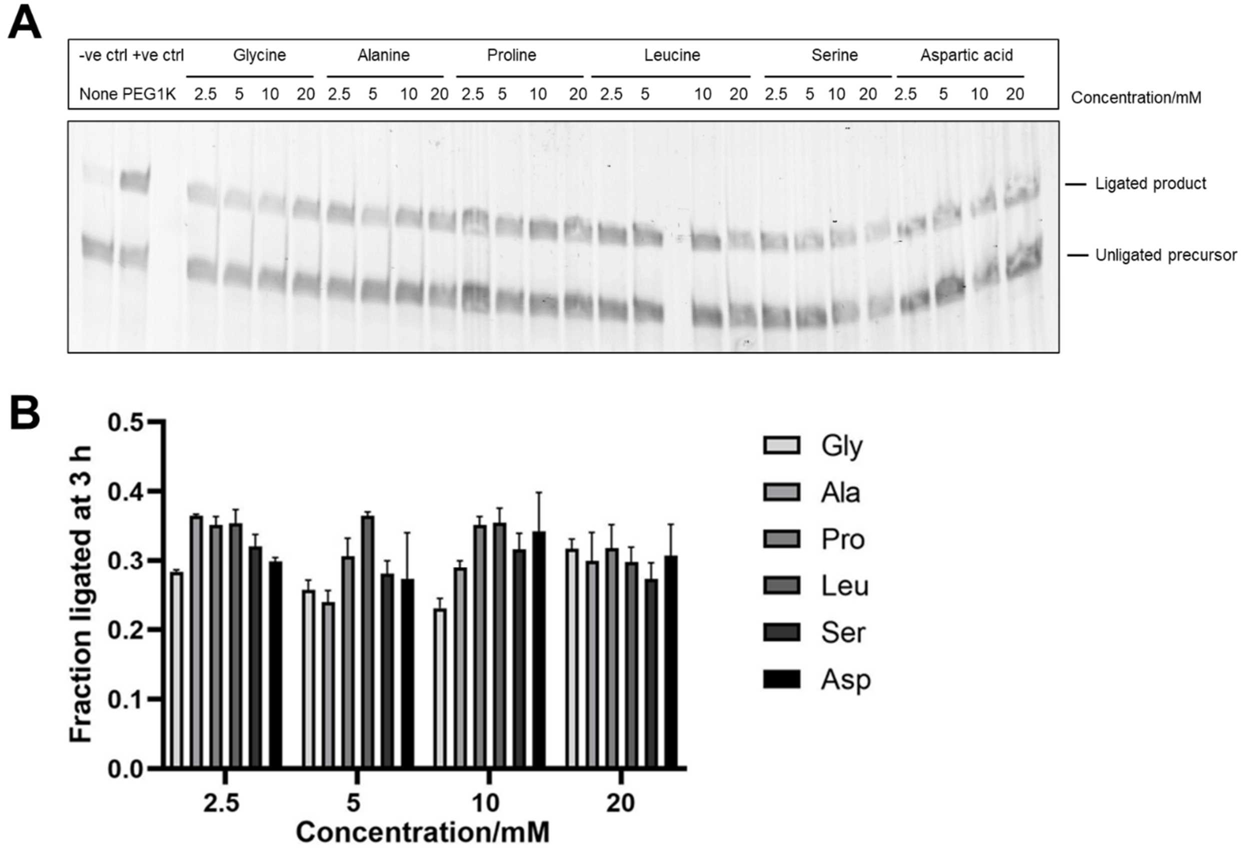


**Figure S10. Ribozyme-catalyzed RNA ligation is stimulated in the presence of prebiotically relevant amino acids. A.** Ligase 1 ribozyme is active at 1 mM $Mg^{2+}$ in the presence of 2.5 mM-20 mM amino acids (Gly, Ala, Pro, Leu, Ser, Asp). 19% PEG 1000 was used as a positive control. **B.** Ligation yields after 3 h were comparable across the different concentrations tested. Ligation reactions contained 1 µM ribozyme, 1.2 µM RNA template, and 2 µM 2-AI-activated RNA substrate in 100 mM Tris-HCl pH 8.0 and 1 mM $MgCl_2$. Reactions were performed in the absence or presence of the indicated amino acids. None indicates the absence of crowders.
